# Intracellular and intercellular gene regulatory networks inference from time-course individual RNA-Seq

**DOI:** 10.1101/2021.05.05.442868

**Authors:** Makoto Kashima, Yuki Shida, Takashi Yamashiro, Hiromi Hirata, Hiroshi Kurosaka

## Abstract

Gene regulatory network (GRN) inference is an effective approach to understand the molecular mechanisms underlying biological events. Generally, GRN inference mainly targets intracellular regulatory relationships such as transcription factors and their associated targets. In multicellular organisms, there are both intracellular and intercellular regulatory mechanisms. Thus, we hypothesize that GRNs inferred from time-course individual (whole embryo) RNA-Seq during development can reveal intercellular regulatory relationships (signaling pathways) underlying the development. Here, we conducted time-course bulk RNA-Seq of individual mouse embryos during early development, followed by pseudo-time analysis and GRN inference. The results demonstrated that GRN inference from RNA-Seq with pseudo-time can be applied for individual bulk RNA-Seq similar to scRNA-Seq. Validation using an experimental-source-based database showed that our approach could significantly infer GRN for all transcription factors in the database. Furthermore, the inferred ligand-related and receptor-related downstream genes were significantly overlapped. Thus, the inferred GRN based on whole organism could include intercellular regulatory relationships, which cannot be inferred from scRNA-Seq based only on gene expression data. Overall, inferring GRN from time-course bulk RNA-Seq is an effective approach for understanding the regulatory relationships underlying biological events in multicellular organisms.

## Introduction

Regulation of gene expression is a fundamental factor that controls cellular events such as proliferation and differentiation. Understanding gene regulatory interconnections is an important step to elucidate the molecular mechanisms underlying cellular events. Recently, gene regulatory network (GRN) inference based on time-course data has attracted much attention in single-cell RNA-Seq (scRNA-Seq). State-of-the-art scRNA-Seq analysis techniques can generate transcriptome information from thousands of cells^1–5^. Transcriptomic heterogeneity of cells due to asynchronous progression of cellular events enables us to infer regulatory relationships of genes. During inference, first, dimensional reduction of scRNA-Seq data provides a trajectory of cellular events such as differentiation and proliferation^6,7^. Then, assignment of pseudo-time can place cells along the trajectory. Since scRNA-Seq with pseudo-time is a dense time-course observation of cellular events, gene regulatory networks can be inferred by comparing the timing of gene upregulation and downregulation along pseudo-time^8,9^.

Originally, GRN inference was applied for gene expression data from tissue and pooled cells (bulk samples), generated using DNA microarray and RNA-Seq^10,11^. Compared to steady-state data, time-course data allows GRN inference by comparing the timing of gene upregulation and downregulation^12–15^. However, GRN inference from time-course data of bulk samples has not been popular due to the following drawbacks: 1) RNA extraction and library preparation of a large number of bulk samples prior to sequencing are too costly and time-consuming^16^, and 2) biological variances may result in inconsistencies between real sampling time and transcriptome status^17^. Fortunately, recent technical advances related to bulk RNA-Seq have overcome these limitations. Advances in sequencing platforms^18^, RNA extraction method^16,19^ and bulk 3′ RNA-Seq library preparation methods^20–22^ have enabled a cost-effective time-course individual RNA-Seq^23^ (a time-series RNA-Seq targeting a whole embryo or tissue of each individual). Pseudo-time analysis for individual RNA-Seq might capture individual differences in progression speed of biological events such as development. Following assignment of pseudo-time to each individual RNA-Seq data, GRN can be inferred similar to an inference based on scRNA-Seq.

Theoretically, GRN inferred from whole-body and tissue RNA-Seq are different from those inferred from scRNA-Seq. scRNA-Seq provides transcriptomic information at the cellular level that enables inference of intracellular GRN involved in the progression of proliferation and differentiation^24^. In contrast, bulk RNA-Seq of whole body and tissues could contain transcriptomic information at the level of cell populations. Time course individual RNA-Seq during development would enable inference of both intracellular and intercellular GRN (cell-cell communications) involved in the developmental process. For instance, during embryonic development, upregulation of ligand-related genes can be followed by upregulation of downstream genes in cells expressing receptor genes^25^.

We thus hypothesize that GRNs inferred from time-course individual RNA-Seq during embryonic development would include intercellular regulatory relationships between ligand genes and downstream genes of related signaling pathways. To test this hypothesis, we conducted time-course bulk RNA-Seq of individual mouse embryos in early development, followed by pseudo-time analysis and GRN inference.

## Results

### Pseudo-time analysis of the time-course bulk RNA-Seq for mouse embryos

To determine whether gene regulatory network (GRN) inference based on time-course bulk RNA-Seq is effective, we conducted time-course bulk RNA-Seq for individual mouse embryos. RNA was extracted from each individual whole embryo (n = 10 or 11) at seven time points: E7.5, E8.5, E9.5, E10.5, E11.5, E12.5, and E.13.5, followed by 3′ RNA-Seq using the ‘Lasy-Seq’ method^21^. As a result, we obtained 76 RNA-Seq datasets with an average of 8.5 million reads per sample. The reads were mapped onto the mouse reference sequence, followed by calculation of the read counts of each gene in each sample. We then used ‘Seurat’ ^26^, an R package used for single cell omics analysis, normalization of read counts, detection of highly variable genes, and dimension reduction of omics date. On the PC1 and PC2 plane obtained with Seurat, samples at the same stage were close to each other (Fig. 1a). As expected, clusters of each stage were ordered according to the developmental process (Fig. 1b). Using an R package ‘slingshot’^27^, we inferred the developmental trajectory of mouse embryos and calculated the pseudo-time for each sample (Fig. 1 a and b). Pseudo-time analysis revealed individual differences among embryos in the speed of their developmental processes. For example, the pseudo-time of a sample at E10.5 and of E11.5 were very close to each other (Fig. 1 a and b). Using the pseudo-time analysis, the time-course RNA-Seq data of the seven time points could be converted into those of 76 time points. Using pseudo-time as temporal information instead of stage (real sampling time) improved the sum of squared residuals (SSR) between observed and fitted values (Fig. 1c and d). The SSRs along pseudo-time were decreased by 0.745% (all genes) and 3.835% (high variable genes), on average, compared with the SSRs along stage. These results indicate that integrating pseudo-time into the analysis, instead of real sampled stage, could improve capture temporal expression dynamics, by considering individual differences in the progression speed of biological events during the mouse early embryonic development.

**Figure 1.**
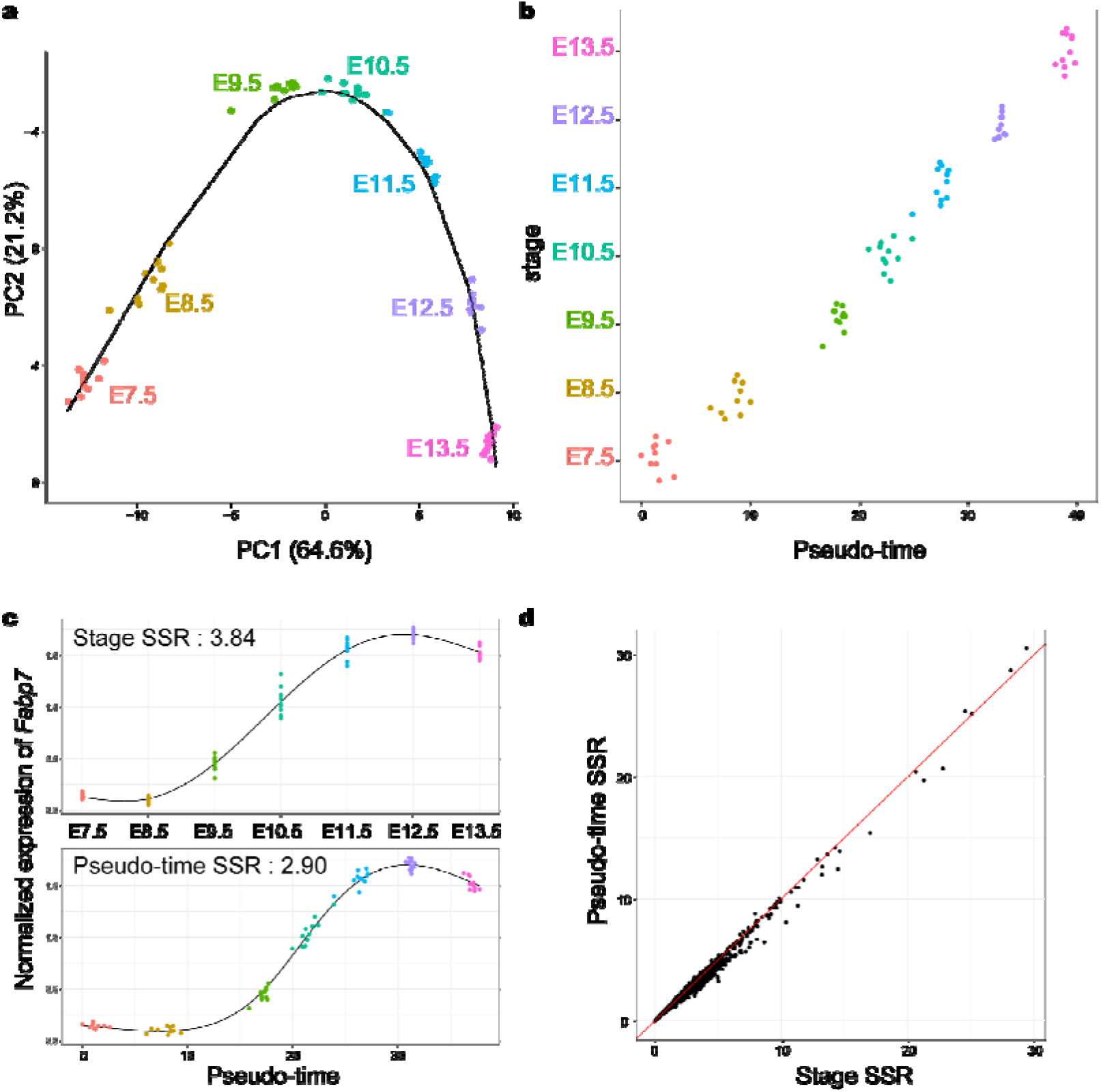
Pseudo-time analysis for time-course individual bulk RNA-Seq of mouse embryos in early development. (a) Transcriptomic trajectory of mouse embryos from E7.5 to E13.5. Each point on the PC1-PC2 plane indicates each individual RNA-Seq result. The solid line indicates an inferred trajectory by slingshot. (b) A scatterplot of pseudo-time and corresponding stage for each sample. Each point indicates each individual RNA-Seq result and is jittered along the Y-axis. (c) An example (*Fabp7*) of difference of gene expression dynamics along stage and pseudo-time. The black lines indicate smoothing curves for each data. Sum of squared residuals (SSR) between observed and fitted values were calculated. (d) A scatterplot of SSR of normalized gene expression of all genes along stage and pseudo-time. The red line indicates the same value between stage and pseudo-time.

### Gene regulatory network inference based on individual RNA-Seq of whole mouse embryos

Next, we inferred a GRN from the dataset of time-course individual RNA-Seq of whole mouse embryos. We used SCODE, which solves linear ordinary differential equations to infer GRN ^8^. To avoid incorrect emphasis on the technical noise of RNA-Seq, we used a non-scaled normalized gene expression matrix as the input for SCODE. Selection of the *D* size, a parameter that affects the number of assumed basic patterns of expression dynamics of the dataset, is important for robust GRN inference, since a needlessly large *D* causes an unstable inferred GRN^8^. In this study, we used *D* = 4 similar to the previous study^8^, with which the SSR was relatively small (Fig. 2a). For 28117 genes whose read counts were greater than zero, SCODE produced *A* (28117 × 28117 matrix) corresponding to the inferred gene regulatory network, in which the value of *A_i,j_* indicates regulatory effects on the downstream gene *i* from the regulator *j*. *A_i,j_* > 0 indicates that the regulator *j* positively regulates gene *i,* whereas *A_i,j_* < 0 indicates the opposite. Because SCODE optimizes *A* by random sampling, we optimized *A* 20 times to check for reproducibility. Then, Pearson’s correlation coefficients between the values of each *A* from the 20 optimizations and the mean*A* were calculated (Fig. 2b). Almost all optimizations produced similar *A* with high correlation coefficients (Fig. 2b). In the subsequent analysis, we used the average values of the top 10 *A* showing higher correlations with the mean*A* from 20 optimizations. We then tried to define the thresholds for significant regulatory relationships between regulators and downstream genes. Because the average expression level of downstream genes showed higher correlation with the absolute values of *A* compared to the regulators (Fig. 2 c and d), we independently defined the threshold for each regulator instead of a constant threshold previously used^8^. For example, absolute values of *A* indicating regulatory relationships between *Sox8* and its regulators showed a positive correlation (Pearson’s correlation coefficient = 0.76) (Fig. 2e), whereas absolute values of *A* showing regulatory relationships between *Sox8* and its downstream genes showed smaller correlations (Pearson’s correlation coefficient = 0.54) (Fig. 2f). Although most of the inferred values of *A* for regulatory relationships between *Sox8* and its downstream genes were around zero, some values were outliers (Fig. 2g). To define the threshold for the downstream genes of *Sox8*, we regressed the linear function for the scatter plot of absolute *A* values for downstream genes in the decreasing order and defined the threshold (Fig. 2g). Finally, in the subsequent analysis, genes with larger absolute values of *A* compared to the threshold were defined as the inferred genes downstream of *Sox8* (Fig. 2g).

**Figure 2.**
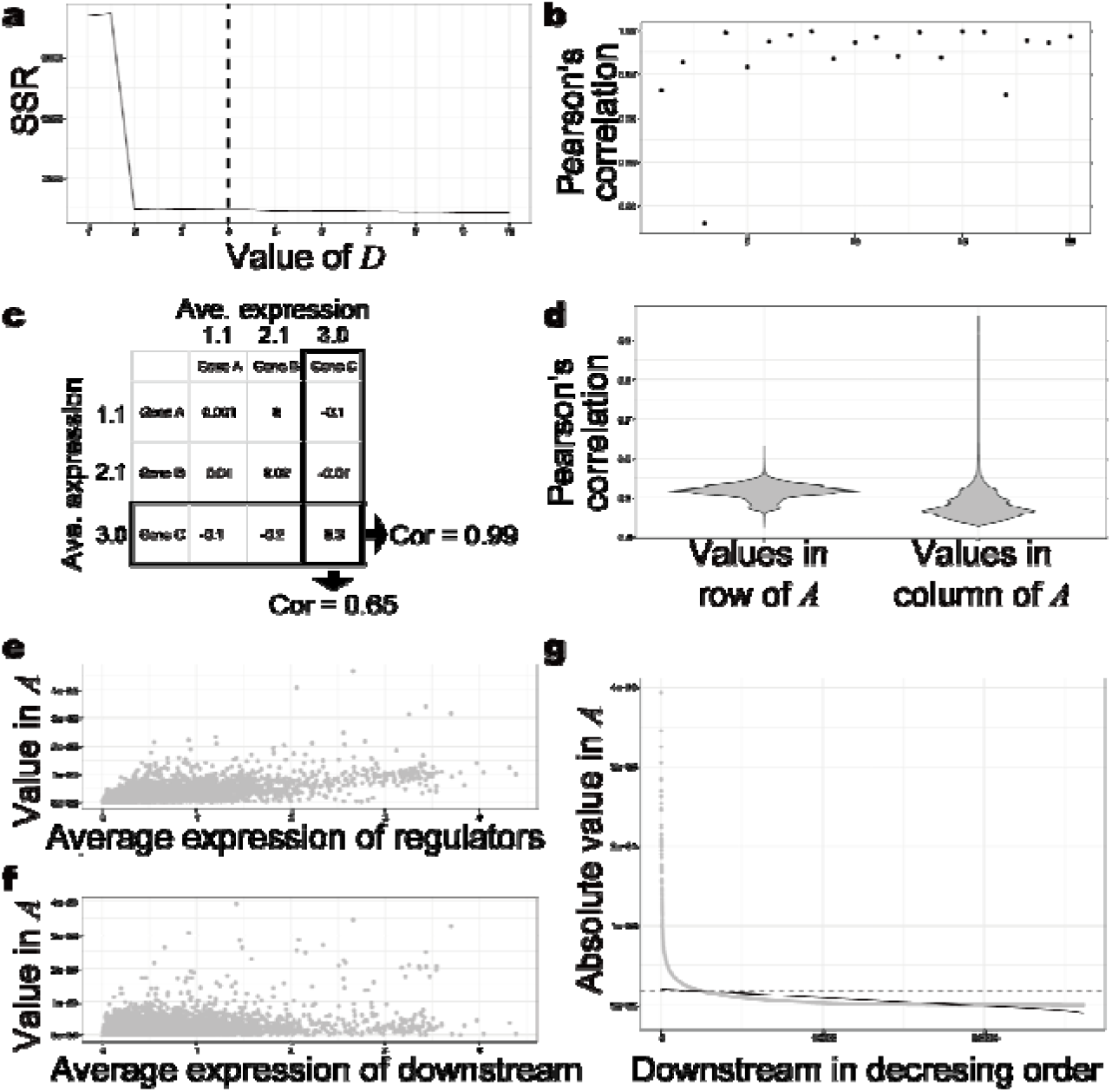
Inference of downstream genes based on the values in *A*. (a) Selection of a parameter *D* for SCODE. The sum of squared residuals (SSR) for each *D* (from 1 to 10 by 0.5) was calculated. We used *D* = 4. (b) Pearson’s correlation coefficients between the values of each *A* from 20 optimizations and the mean*A,* which is the average of each value from all optimizations. (c) Calculation of Pearson’s correlations between the average expression level of each gene and the absolute values in each row and column of *A*. (d) The violin plots of Pearson’s correlations (e, f) Scatter plot of the average expression level of each gene and the absolute values in the column (e) and row (f) of *A* for *Sox8*. (g) An example of the definition of thresholds for significant regulatory relationship between the regulator and downstream genes. A scatter plot of the values of *A* for a regulator, *Sox8,* in decreasing order. The solid line indicates a regression for the scatter plot. The dashed lines indicate the defined thresholds.

### Validation of the inferred network

Next, we evaluated the inferred GRN by comparing the inferred genes downstream of the TFs with the information in the TF2DNA database, which is an experimental-source-based database of the binding motifs and downstream genes for 438 mouse TFs^28–39^. First, we evaluated the effectiveness of target prediction based on the absolute values of *A* by calculating the area under the curve (AUC) (Fig. 3a), obtaining an average AUC of 0.704, suggesting that the inferred regulatory relationships could reflect the true regulatory network. Also, we calculated AUC for inferred regulatory relationships using dynGENIE3^40^, an algorithm using sampled stage information as temporal information to infer GRN. The average AUC was 0.50 (Fig. 3a), suggesting that advantage of usage of pseudo-time in GRN inference based on time-course individual RNA-Seq. Second, we examined the validity of the defined thresholds by assessing the differences between the validated target gene rate (number of validated target genes / number of all inferred downstream genes) above the defined thresholds and the best validated rate (Fig. 3b). Below the threshold, validated target gene rates for the 438 TFs were 0.69% smaller than the best validated target gene rates on average. The validated target genes were 67.5% of the inferred downstream genes on average (Table S1, Fig. 3c and d). Compared with the background rate (number of known target genes / number of all genes), the validated target gene rates of the inferred downstream genes of all TFs were statistically higher (adjusted p-value < 0.01) (Fig. 3d). These results suggest that our approach could infer the GRN underlying early development in mice.

**Figure 3.**
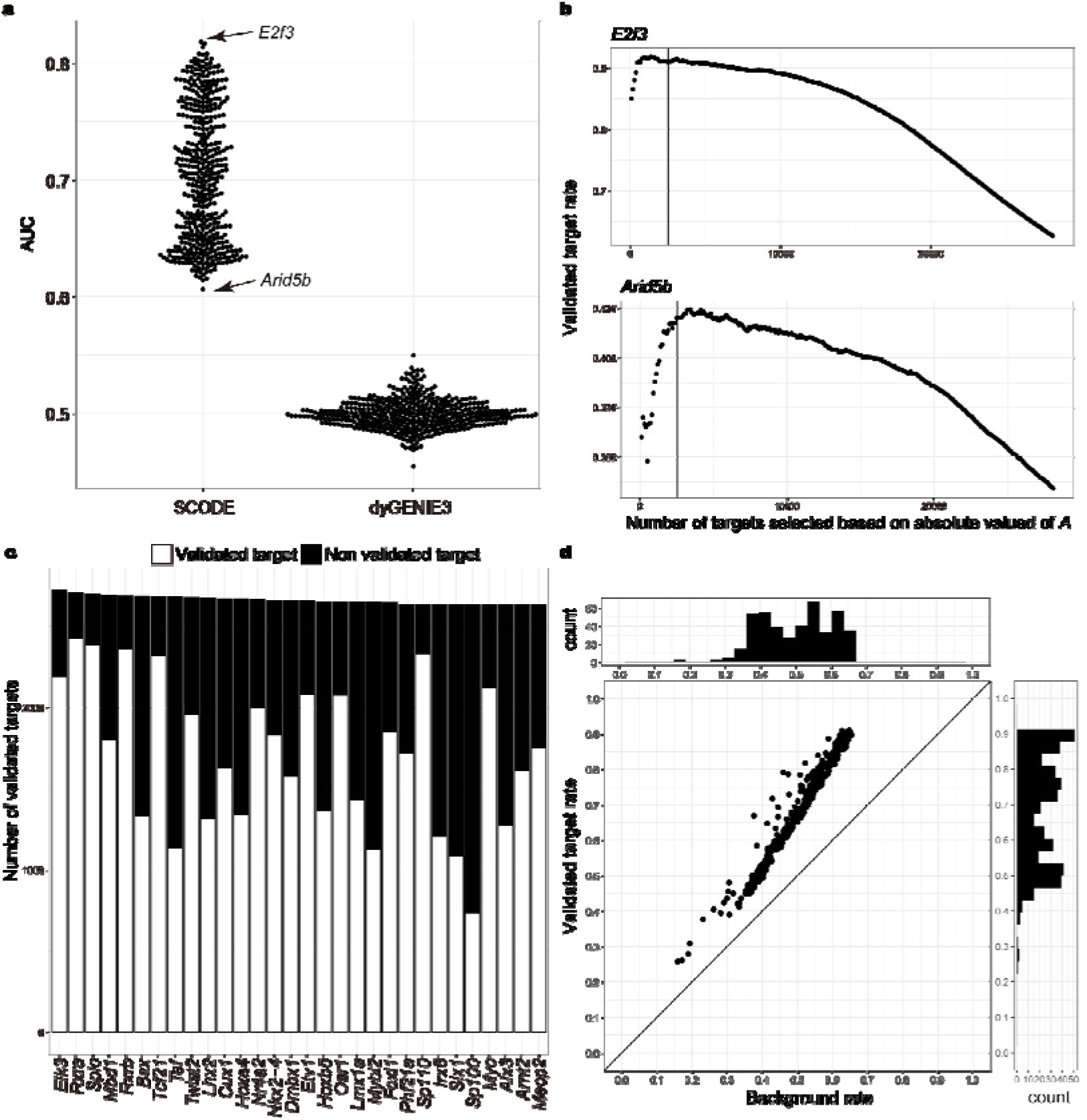
Validation of the inferred regulatory network of transcription factors with the TF2DNA database. (a) Scatter plot of area under the curve (AUC) target selection for each transcription factor (TF) based on the absolute values of *A* inferred using SCODE and weight.matrix inferred using dyGENIE3. (b) Scatter plots of the validated inferred target gene rate of *E2f3* and *Arid5b* in the TF2DNA database. Downstream genes of TFs were selected based on the absolute values of *A* in decreasing order. Solid lines indicate genes at the thresholds. (c) Bar graph of validated and non-validated downstream genes of each TF in the TF2DNA database, in decreasing order of the total number of inferred downstream genes. Only the top 30 TFs are shown. (d) Scatter plot of the validated target gene rate of the inferred downstream genes of TF*j* and the background rate of target genes in the TF2DNA database. Histograms show the distribution of validated target gene rates of inferred downstream genes and the background rates of downstream genes in the TF2DNA database.

### The inferred network contained regulatory relationships involved in cell-cell interaction

Our GRN inference was based on bulk RNA-Seq containing the information of all cells in the body. We thus hypothesized that the inferred GRN also included the intercellular regulatory network. To examine this possibility, we checked the overlaps of inferred downstream genes for genes related to the ligands and receptors of nine major signaling pathways (Fig. 4a): Wnt/βcatenin, TNF, TGF-β, Hedgehog, FGF, EGF, Delta/Notch, BMP, and Retinoic acid (RA) signaling pathways (Table S2). As expected, the inferred downstream genes of all pairs of ligand and receptor genes were significantly overlapped (adjusted p-value < 0.01) (Fig. 4a). On average, 94.2% of the inferred downstream genes for the ligand-related and receptor-related genes were overlapped (Fig. 4A). For example, 4510 genes were inferred as the downstream of *Wnt* genes, and 4545 genes were downstream of *Fzd* genes. However, 97.5% of the inferred downstream of *Wnt* genes were also inferred as the downstream of *Fzd* genes (Fig. 4a). Since SCODE can infer whether each downstream gene is positively or negatively regulated^8^, we assessed the overlaps of positively and negatively regulated gene downstream of the representative ligand-related and receptor-related genes of each signaling pathway, i.e., with the largest number of inferred downstream genes among each gene family (Fig. 4 b-d and Supplementary Figure 1,2). For six signaling pathways (TNFβ, TGF, FGF, EGF, Delta/Notch, and BMP signaling pathways), which activate the downstream genes only when ligands bind to the receptors^41^, the regulatory directions of ligand-related and receptor-related genes for most inferred downstream genes were the same (Fig. 4b and Supplementary Figure 1). In contrast, in case of the RA signaling pathway, wherein RA receptors function as transcriptional repressors without RA binding (Supplementary Figure 2a)^42^, the regulatory directions of a ligand-related gene, *Aldh1a3,* which encodes a protein involved in RA synthesis, and a receptor-related gene, *Rara,* for the downstream genes were opposite (Fig. 4c). In case of the Hedgehog signaling pathway, the regulatory directions of *Ssh2* and *Ptchd4* tended to be opposite (Fig. 4d). The regulatory directions of *Ssh2* and *Gli3* for the downstream genes were also opposite (Fig. 4d). In the absence of Hedgehog ligands, the full-length Gli family proteins are ubiquitinated and function as repressors (Supplementary Figure 2b)^43^. With the binding of Hedgehog ligands, PATCHED inhibits the degradation of Gli family proteins and allows them to function as transcriptional activators (Supplementary Figure 2a)^43^. Thus, the overlap of downstream genes that are positively and negatively regulated by *Ssh2, Ptchd4,* and *Gli3* is reasonable. In case of the Wnt/βcatenin signaling pathway, the regulatory directions of *Wnt7a* and *Fzd5* for the downstream were opposite (Supplementary Figure 3a). Our time-course RNA-Seq revealed that among the *Fzd* and *Wnt* genes, *Fzd5* and *Wnt8a,* but not *Wnt7a,* were the only genes expressed dominantly in the early developmental stage among the protein families. (Supplementary Figures 4 and 5). Similar regulatory directions were found for most of the inferred downstream genes of *Wnt8a-Fzd5*, which could be a functional pair in the early developmental stages. Furthermore, the regulatory directions for most of the inferred downstream genes of *Wnt7a-Fzd6* (*Fzd* genes with the second most inferred downstream genes) were also the same (Supplementary Figure 3). In conclusion, our approach allowed successful inference the intercellular regulatory relationships related to the major signaling pathways as well as the intracellular pathways related to TFs.

**Figure 4.**
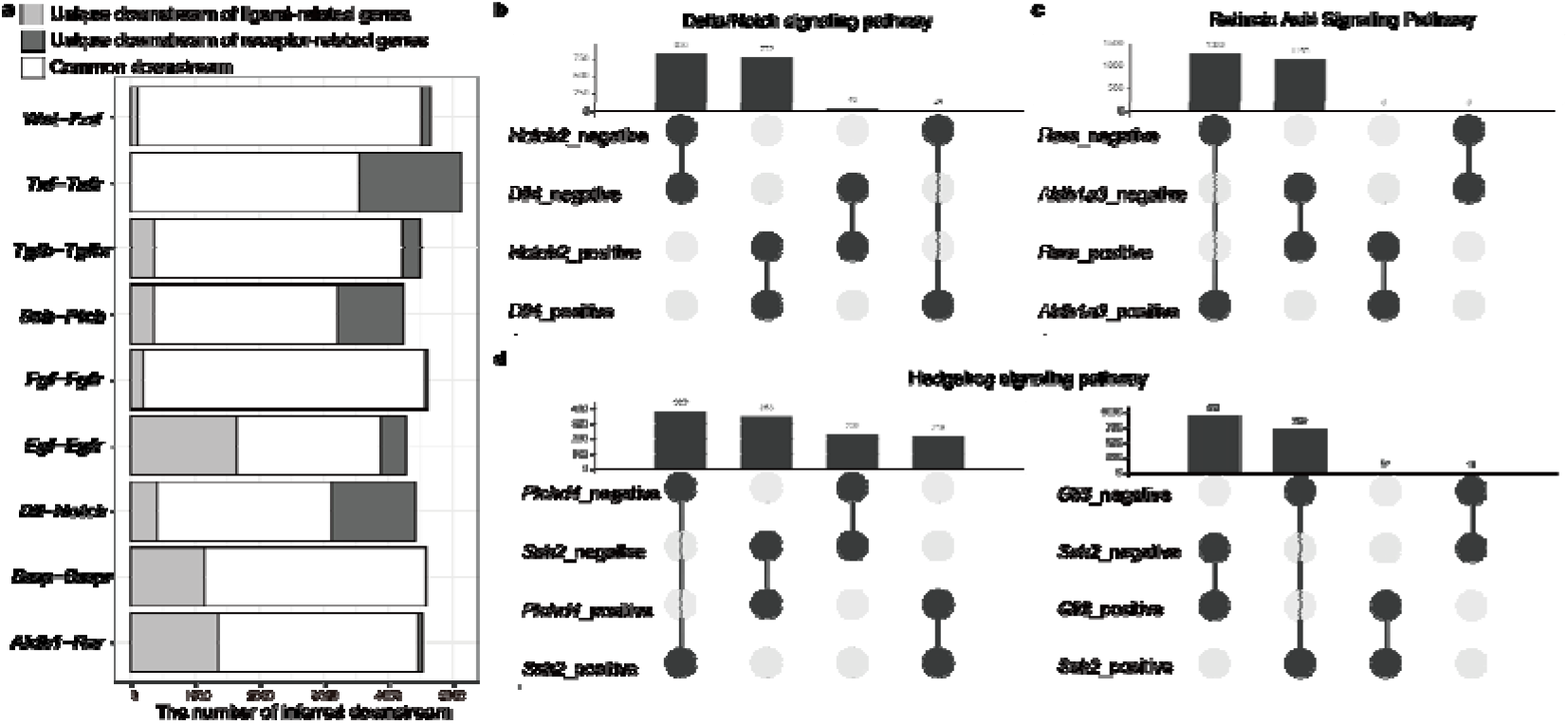
Overlap of inferred genes downstream of ligand-related and receptor-related genes. (a) Bar plot of the number of inferred downstream genes that are common and unique for each ligand-receptor pair. (b-d) Upset plots of inferred downstream genes that are positively and negatively regulated by the representative ligand-related and receptor-related genes. (b) Retinoic acid signaling pathway. (c) Delta/Notch signaling pathway. (d) Hedgehog signaling pathway.

## Discussion

Recently, GRN inference based on the combination of scRNA-Seq and pseudo-time analysis has attracted much attention^44^. However, to our knowledge, this study is the first to report GRN inference based on the combination of individual bulk RNA-Seq and pseudo-time analysis. Herein, pseudo-time explained variances of gene expression during mouse early development better than real sampled stage (Fig. 1 c and d), suggesting that the variance of gene expression observed using our time-course individual RNA-Seq was not mere stochastic variability in gene expression but individual difference in the progression speed of development. Thus, pseudo-time uses the correlation of gene expression dynamics more effectively than the real temporal information in time-course individual RNA-Seq.

Unlike scRNA-Seq, which can provide transcriptomic dynamics in a certain cellular event such as proliferation and differentiation, bulk RNA-Seq could provide a mixture of various transcriptomic dynamics regarding the cellular events occurring in an embryo^45^. Theoretically, GRN inference based on bulk RNA-Seq cannot provide multiple lineage-specific regulatory relationships. This may explain why GRN inference based on bulk RNA-Seq has not been attempted as with scRNA-Seq. In this study, we succeeded in inferring the known regulatory relationships of the TFs in the TF2DNA database with high AUC compared to the GRN inferred from scRNA-Seq^46^ (Fig. 3). This suggests that GRN inference from bulk RNA-Seq provides insights into the regulatory interconnections of genes, even though GRN inferred from bulk RNA-Seq would have limitations in the comprehensiveness of inferred regulatory relationships compared to those from scRNA-Seq. Furthermore, drastic changes in the cell population occur during early developments. It is possible that our GRN inferred from bulk RNA-Seq merely reflect the changes in cell populations but not the interconnections of genes. Considering the results of the inference for *Wnt* and *Fzd* genes (Supplementary Figure 2), these points should be considered when interpreting the inferred GRN based on the time-course bulk RNA-Seq.

Assignment of pseudo-time is an important step in GRN inference from time-course data. In case of scRNA-Seq, the accuracy of pseudo-time assignment is controversial^47^. In contrast, in the case of time-course individual bulk RNA-Seq, the accuracy of pseudo-time assignment could be assured by the actual sampling time, which can be an advantage compared to GRN inference from scRNA-Seq.

In this study, we proposed a new strategy for threshold of significant gene regulatory relationships inferred by SCODE (Fig. 2). In the case of *E2f3,* the number of predicted target genes is only ~10% of the ground truth targets under our strategy. Shifting the threshold to approximately 17,000 targets for *E2f3* may still result in a validated target rate of approximately 80% (Fig. 3b). Since decreasing threshold may produce false positive inferred relationships for some genes, the determination of the threshold remains challenging. Considering high AUCs for the TFs in the database (Fig. 3a), the threshold should change depending on the study aims.

GRN indicates the intracellular interconnections of genes in a narrow sense; intercellular regulation of genes via cell-cell communication is also a key factor to understand the regulatory mechanisms underlying multicellular organisms. Several studies have attempted to systematically identify cell-cell communications based on single cell gene expression profiles and information regarding ligand-receptor pairs^48–51^. Since these approaches required prior knowledge, they could only be applied for the major model organisms and could not reveal novel signaling pathways. We demonstrated that GRN inference based on time-course individual RNA-Seq could infer intercellular regulatory relationships related to cell-cell communication via cell signaling pathways. This approach only requires time-course RNA-Seq results and is applicable for non-model organisms without ligand-receptor information. Theoretically, the GRN inferred based on scRNA-Seq cannot include intercellular interconnections of genes. Comparison of GRN from scRNA-Seq and bulk RNA-Seq would be useful for systematic identification of genes involved in cell-cell communications during cellular events of interest. Taken together, our approach is a powerful tool for understanding intracellular and intercellular regulatory relationships of genes, which cannot be achieved using the existing GRN inferences based on scRNA-Seq alone. As discussed above, bulk RNA-Seq is limited in the comprehensiveness of inferred regulatory relationship since multiple lineage-specific regulatory relationships cannot be deduced from GRN inference based on bulk RNA-Seq. A future novel bioinformatic approach that can deconvolute gene expression in each tissue and cell lineage from bulk RNA-Seq will help overcoming this limitation.

## Methods

### Maintenance of mice

Embryos were collected from pregnant female Institute of Cancer Research (ICR) mice (CLEA, Tokyo, Japan) at each stage (E7.5, E8.5, E9.5, E10.5, E11.5, E12.5, and E.13.5). The number of replicates (embryos) were 10 at E7.5 and 11 at the remaining stages. All sacrificed female mice were housed under a 12 h dark-light cycle in which the light phase started from 8 am. All animal experiments were performed in accordance with the guidelines of the Animal Care and Use Committee of Osaka University Graduate School of Dentistry, Osaka, Japan.

### RNA extraction

Total RNA extraction was performed using the RNeasy® kit (QIAGEN, Hilden, Germany) according to manufacturer’s protocol. The total RNA concentration was measured using Qubit™ RNA HS Assay Kit (Thermo Fisher Scientific, Waltham, MA, USA) and was adjusted to 5 ng/ul and stored at −80 □ until subsequent analysis.

### RNA-Seq library preparation and sequencing

Non targeted RNA-Seq were conducted according to the Lasy-Seq ver. 1.1 protocol (https://sites.google.com/view/lasy-seq/)^21,23^. Briefly, 50 ng of total RNA were reverse transcribed using an RT primer with index and SuperScript IV reverse transcriptase (Thermo Fisher Scientific, Waltham, MA, USA). Then, all RT mixtures of the samples were pooled and purified using an equal volume of AMpure XP beads (Beckman Coulter, Brea, CA, USA) according to the manufacturer’s instructions. Second strand synthesis was conducted on the pooled samples using RNaseH (5 U/μL, Enzymatics, Beverly, MA, USA), and DNA polymerase I (10 U/μL, Enzymatics, Beverly, MA, USA). To avoid the carryover of large amounts of rRNAs, the mixture was subjected to RNase treatment using RNase T1 (Thermo Fisher Scientific, Waltham, MA, USA). Then, purification was conducted with a 0.8× volume of AMpure XP beads. Fragmentation, end-repair, and A-tailing were conducted using 5× WGS Fragmentation Mix (Enzymatics, Beverly, MA, USA). The Adapter for Lasy-Seq was ligated using 5× Ligation Mix (Enzymatics, Beverly, MA, USA), and the adapter-ligated DNA was purified twice with a 0.8× volume of AMpure XP beads. After optimisation of PCR cycles for library amplification by qPCR using EvaGreen, 20× in water (Biotium, Fremont, CA, USA) and the QuantStudio5 Real-Time PCR System (Applied Biosystems, Waltham, MA, USA), the library was amplified using KAPA HiFi HotStart ReadyMix (KAPA BIOSYSTEMS, Wilmington, MA, USA) on the ProFlex PCR System (Applied Biosystems, Waltham, MA, USA). The amplified library was purified with an equal volume of AMpure XP beads. One microliter of the library was then used for electrophoresis using a Bioanalyzer 2100 with the Agilent High Sensitivity DNA kit (Agilent Technologies, Santa Clara, CA, USA) to assess quality. Then, sequencing of 150-bp paired-end reads was performed using HiSeq X Ten (Illumina, San Diego, CA, USA).

### Mapping and gene quantification

Read 1 reads were processed with fastp (version 0.21.0)^52^ using the following parameters: --trim_poly_x -w 20 --adapter_sequence=AGATCGGAAGAGCACACGTCTGAACTCCAGTCA

--adapter_sequence_r2=AGATCGGAAGAGCGTCGTGTAGGGAAAGAGTGT -l 31. The trimmed reads were then mapped to the mouse reference sequences of Mus_musculus.GRCm38.cdna.all.fa, using BWA mem (version 0.7.17-r1188)^53^ with the default parameters. The read count for each gene was calculated with salmon using -l IU, which specifies the library type (version v0.12.0)^54^. Then, using R (version 4.0.1)^55^, the sums of read counts per gene were calculated. Genes with read counts greater than zero were used in the subsequent analysis.

### Pseudo-time analysis

Read counts were normalized using the “NormalizeData” function with the default parameters in Seurat (version 4.0.0)^56^, which produces natural-log transformed (read per 10,000 + 1). For principal component analysis (PCA), the normalized read counts were centered but not scaled using the “ScaleData” function with the default parameters except for do.scale = F. Then, PCA was conducted using the “RunPCA” function for genes with high dispersion, which were selected using the “FindVariableFeatures” function with default parameters except for selection.method = “mvp”. Finally, SingleCellExperiment (version 1.10.1)^57^ and slingshot (version 1.6.1)^27^ were used to calculate the pseudo-time for each sample.

### Evaluation of considering pseudo-time instead of stage

The “smooth.spline” function in R (version 4.0.1)^55^ with the default parameters except for “all.knots = T, lambda = 0.001” was used to obtain smoothed curve for normalized expression of each gene in the “data” slot of the Seurat object, and stage or pseudo-time. Then sum of squared residuals (SSR) between observed and fitted values was calculated for each gene. Mean of SSRs were calculated against all genes and the high variable genes obtained with the “FindVariableFeatures” function.

### Gene regulatory network inference

In the SCODE algorithm^8^, normalized expression data in the “data” slot of the Seurat object, and pseudo-time were used to infer GRN. *A* was optimized 20 times with one hundred iterations and *D = 4*. Pearson’s correlation coefficients between values of each *A* from the 20 optimizations and the mean*A* (the average of each value of *A*) were calculated. In the following analysis, we used the average values of the top 10 *A* showing higher correlations with the mean*A* from 20 optimizations. To define the thresholds for downstream gene selection for each gene, the linear function was regressed using the “nls” function in R for the scatter plot of absolute values of *A* for downstream genes in the decreasing order; the X and Y axes showed the integers from 1 to 28,117, and the absolute values of *A*, respectively. Genes with larger absolute values of *A* compared to the Y values of the regressed line were defined as the inferred gene downstream of each gene.

In the dynGENIE3 algorithm^40^, normalized expression data in the “data” slot of the Seurat object, and samples stage were used to infer GRN. E7.5, E8.5, E9.5, E10.5, E.11.5, E12.5, and E.13.5 were converted into 1, 2, 3, 4, 5, 6, and 7, respectively. The all parameters were used with the default values. weight.matrix inferred by dynGENIE3 was used as inferred regulatory relationships.

### Evaluation of inferred GRN by comparing with a TF-downstream gene database

For validating the inferred GRN, information regarding the binding motifs and targets for 438 mouse TFs in the TF2DNA database was used^28–39^. AUC values for downstream prediction based on the absolute values of *A* were calculated using the “performance” function in ROCR (version 1.0-11)^58^. Statistical analysis for the enrichment of validated target genes among the inferred genes was conducted using the “fisher.test” function in R. Statistical analysis for the overlap of inferred downstream genes for ligand-related and receptor-related genes was conducted using the “enrichment_test” function in Rvenn (version 1.1.0)^59^. For all statistical tests, BH correction was performed using the “p.adjust” function. The upset plots were drawn using the “upset” function in UpSetR (version 1.4.0)^60^.

## Acknowledgments

This work was supported by Japan Society for the Promotion of Science [grant number JP19H03858 to H.K.]. We would like to thank Editage (www.editage.com) for English language editing.

## Author contributions

M.K.,Y.S., and H.K. conceived and conducted the experiments, and M.K. analyzed the results and wrote the manuscript. All authors reviewed and approved the final manuscript.

## Data availability

All RNA-Seq sequence data is deposited in PRJNA725414 of SRA. The R source code used in the present study is available at https://github.com/Makoto-Kashima/TIODE

## References

1. Sasagawa, Y. et al. Quartz-Seq2: A high-throughput single-cell RNA-sequencing method that effectively uses limited sequence reads. Genome Biol. 19, 1–24 (2018).

2. Sasagawa, Y. et al. Quartz-Seq: a highly reproducible and sensitive single-cell RNA-Seq reveals non-genetic gene expression heterogeneity. Genome Biol. 14, R31 (2013).

3. Gao, C., Zhang, M. & Chen, L. The Comparison of Two Single-cell Sequencing Platforms: BD Rhapsody and 10x Genomics Chromium. Curr. Genomics 21, 602–609 (2020).

4. Hayashi, T. et al. Single-cell full-length total RNA sequencing uncovers dynamics of recursive splicing and enhancer RNAs. Nat. Commun. 9, 1–16 (2018).

5. Klein, A. M. et al. Droplet barcoding for single-cell transcriptomics applied to embryonic stem cells. Cell 161, 1187–1201 (2015).

6. Treutlein, B. et al. Reconstructing lineage hierarchies of the distal lung epithelium using single-cell RNA-seq. Nature 509, 371–375 (2014).

7. Haghverdi, L., Büttner, M., Wolf, F. A., Buettner, F. & Theis, F. J. Diffusion pseudotime robustly reconstructs lineage branching. Nat. Methods 13, 845–848 (2016).

8. Matsumoto, H. et al. SCODE: an efficient regulatory network inference algorithm from single-cell RNA-Seq during differentiation. Bioinformatics 33, 2314–2321 (2017).

9. Aalto, A., Viitasaari, L., Ilmonen, P., Mombaerts, L. & Gonçalves, J. Gene regulatory network inference from sparsely sampled noisy data. Nat. Commun. 11, 3493 (2020).

10. Ko, D. K. & Brandizzi, F. Network-based approaches for understanding gene regulation and function in plants. Plant Journal vol. 104 302–317 (2020).

11. Fernandez-Valverde, S. L., Aguilera, F. & Ramos-Dıaz, R. A. Inference of developmental gene regulatory networks beyond classical model systems: New approaches in the post-genomic era. Integr. Comp. Biol. 58, 640–653 (2018).

12. Iglesias-Martinez, L. F., Kolch, W. & Santra, T. BGRMI: A method for inferring gene regulatory networks from time-course gene expression data and its application in breast cancer research. Sci. Rep. 6, 1–12 (2016).

13. Zhang, J. et al. Differential regulatory network-based quantification and prioritization of key genes underlying cancer drug resistance based on time-course RNA-seq data. PLOS Comput. Biol. 15, e1007435 (2019).

14. Krouk, G., Mirowski, P., LeCun, Y., Shasha, D. E. & Coruzzi, G. M. Predictive network modeling of the high-resolution dynamic plant transcriptome in response to nitrate. Genome Biol. 11, R123 (2010).

15. Ogami, K. et al. Computational gene network analysis reveals TNF-induced angiogenesis. BMC Syst. Biol. 6, 3–8 (2012).

16. Yoshino, K., Nishijima, R. & Kawakatsu, T. Low-cost RNA extraction method for highly scalable transcriptome studies. Breed. Sci. (2020) doi:10.1270/jsbbs.19170.

17. Kilfoil, M. L., Lasko, P. & Abouheif, E. Stochastic variation: From single cells to superorganisms. HFSP J. 3, 379–385 (2009).

18. Muir, P. et al. The real cost of sequencing: scaling computation to keep pace with data generation. Genome Biol. 17, 53 (2016).

19. Ujibe, K., Nishimura, K., Kashima, M. & Hirata, H. Direct-TRI: High-throughput RNA-extracting method for all stages of zebrafish development. BIO-PROTOCOL (2021).

20. Alpern, D. et al. BRB-seq: ultra-affordable high-throughput transcriptomics enabled by bulk RNA barcoding and sequencing. Genome Biol. 20, 71 (2019).

21. Kamitani, M., Kashima, M., Tezuka, A. & Nagano, A. J. Lasy-Seq: a high-throughput library preparation method for RNA-Seq and its application in the analysis of plant responses to fluctuating temperatures. Sci. Rep. 9, 7091 (2019).

22. Li, Y. et al. Decode-seq: A practical approach to improve differential gene expression analysis. Genome Biol. 21, 66 (2020).

23. Kashima, M., Kamitani, M., Nomura, Y., Hirata, H. & Nagano, A. J. DeLTa-Seq: direct-lysate targeted RNA-Seq from crude tissue lysate. (8/15 words) Running Title: Development of direct-lysate targeted RNA-Seq method Corresponding Author. bioRxiv 2020.09.15.299180 (2020) doi:10.1101/2020.09.15.299180.

24. Lam, K. Y., Westrick, Z. M., Müller, C. L., Christiaen, L. & Bonneau, R. Fused Regression for Multi-source Gene Regulatory Network Inference. PLoS Comput. Biol. 12, 1–23 (2016).

25. Basson, M. A. Signaling in Cell Differentiation and Morphogenesis. Cold Spring Harb. Perspect. Biol. 4, 1–21 (2012).

26. Stuart, T. et al. Comprehensive Integration of Single-Cell Data. Cell 177, 1888–1902.e21 (2019).

27. Street, K. et al. Slingshot: Cell lineage and pseudotime inference for single-cell transcriptomics. BMC Genomics 19, 477 (2018).

28. Pujato, M., Kieken, F., Skiles, A. A., Tapinos, N. & Fiser, A. Prediction of DNA binding motifs from 3D models of transcription factors; identifying TLX3 regulated genes. Nucleic Acids Res. 42, 13500–13512 (2014).

29. Badis, G. et al. Diversity and complexity in DNA recognition by transcription factors. Science (80-.). 324, 1720–1723 (2009).

30. Wei, G. H. et al. Genome-wide analysis of ETS-family DNA-binding in vitro and in vivo. EMBO J. 29, 2147–2160 (2010).

31. Weirauch, M. T. et al. Evaluation of methods for modeling transcription factor sequence specificity. Nat. Biotechnol. 31, 126–134 (2013).

32. Berger, M. F. et al. Compact, universal DNA microarrays to comprehensively determine transcription-factor binding site specificities. Nat. Biotechnol. 24, 1429–1435 (2006).

33. Mathelier, A. et al. JASPAR 2014: An extensively expanded and updated open-access database of transcription factor binding profiles. Nucleic Acids Res. 42, D142–D147 (2014).

34. Matys, V. et al. TRANSFAC and its module TRANSCompel: transcriptional gene regulation in eukaryotes. Nucleic Acids Res. 34, D108 (2006).

35. Weirauch, M. T. et al. Determination and inference of eukaryotic transcription factor sequence specificity. Cell 158, 1431–1443 (2014).

36. Berger, M. F. et al. Variation in Homeodomain DNA Binding Revealed by High-Resolution Analysis of Sequence Preferences. Cell 133, 1266–1276 (2008).

37. Chen, L., Wu, G. & Ji, H. hmChIP: a database and web server for exploring publicly available human and mouse ChIP-seq and ChIP-chip data. Bioinformatics 27, 1447–1448 (2011).

38. Jolma, A. et al. DNA-binding specificities of human transcription factors. Cell 152, 327–339 (2013).

39. Sebé-Pedrós, A. et al. Early eèolution of the T-box transcription factor family. Proc. Natl. Acad. Sci. U. S. A. 110, 16050–16055 (2013).

40. Huynh-Thu, V. A. & Geurts, P. DynGENIE3: Dynamical GENIE3 for the inference of gene networks from time series expression data. Sci. Rep. 8, (2018).

41. Gilbert, S. F. & Barresi, M. J. F. DEVELOPMENTAL BIOLOGY, 11TH EDITION 2016. Am. J. Med. Genet. Part A 173, 1430–1430 (2017).

42. Glass, C. K. & Rosenfeld, M. G. The coregulator exchange in transcriptional functions of nuclear receptors. Genes and Development vol. 14 121–141 (2000).

43. Skoda, A. M. et al. The role of the hedgehog signaling pathway in cancer: A comprehensive review. Bosnian Journal of Basic Medical Sciences vol. 18 8–20 (2018).

44. Dai, H., Jin, Q. Q., Li, L. & Chen, L. N. Reconstructing gene regulatory networks in single-cell transcriptomic data analysis. Zool. Res. 41, 599–604 (2020).

45. Chasman, D. & Roy, S. Inference of cell type specific regulatory networks on mammalian lineages. Current Opinion in Systems Biology vol. 2 130–139 (2017).

46. Chen, S. & Mar, J. C. Evaluating methods of inferring gene regulatory networks highlights their lack of performance for single cell gene expression data. BMC Bioinformatics 19, 232 (2018).

47. Tritschler, S. et al. Concepts and limitations for learning developmental trajectories from single cell genomics. Development (Cambridge) vol. 146 (2019).

48. Kumar, M. P. et al. Analysis of Single-Cell RNA-Seq Identifies Cell-Cell Communication Associated with Tumor Characteristics. Cell Rep. 25, 1458–1468.e4 (2018).

49. Jin, S. et al. Inference and analysis of cell-cell communication using CellChat. Nat. Commun. 12, 1–20 (2021).

50. Cabello-Aguilar, S. et al. SingleCellSignalR: Inference of intercellular networks from single-cell transcriptomics. Nucleic Acids Res. 48, 55 (2021).

51. Wang, S., Karikomi, M., Maclean, A. L. & Nie, Q. Cell lineage and communication network inference via optimization for single-cell transcriptomics. Nucleic Acids Res. 47, 66 (2019).

52. Chen, S., Zhou, Y., Chen, Y. & Gu, J. fastp: an ultra-fast all-in-one FASTQ preprocessor. Bioinformatics 34, i884–i890 (2018).

53. Li, H. & Durbin, R. Fast and accurate short read alignment with Burrows-Wheeler transform. Bioinformatics 25, 1754–1760 (2009).

54. Patro, R., Duggal, G., Love, M. I., Irizarry, R. A. & Kingsford, C. Salmon provides fast and bias-aware quantification of transcript expression. Nat. Methods 14, 417–419 (2017).

55. R Core Team. R: A language and environment for statistical computing. R Foundation for Statis-tical Computing, Vienna, Austria. https://www.r-project.org/ (2017).

56. Hao, Y. et al. Integrated analysis of multimodal single-cell data. bioRxiv 2020.10.12.335331 (2020) doi:10.1101/2020.10.12.335331.

57. Amezquita, R. A. et al. Orchestrating single-cell analysis with Bioconductor. Nat. Methods 17, 137–145 (2020).

58. Sing, T., Sander, O., Beerenwinkel, N. & Lengauer, T. ROCR: visualizing classifier performance in R. Bioinformatics 21, 3940–3941 (2005).

59. Akyol, T. Y. RVenn: An R package for set operations on multiple sets. https://cran.r-project.org/web/packages/RVenn/vignettes/vignette.html (2019).

60. Conway, J. R., Lex, A. & Gehlenborg, N. UpSetR: an R package for the visualization of intersecting sets and their properties. Bioinformatics 33, 2938–2940 (2017).

