## supplemental_figure for "Intracellular and intercellular gene regulatory networks inference from time-course individual RNA-Seq"

**Supplementary Data**

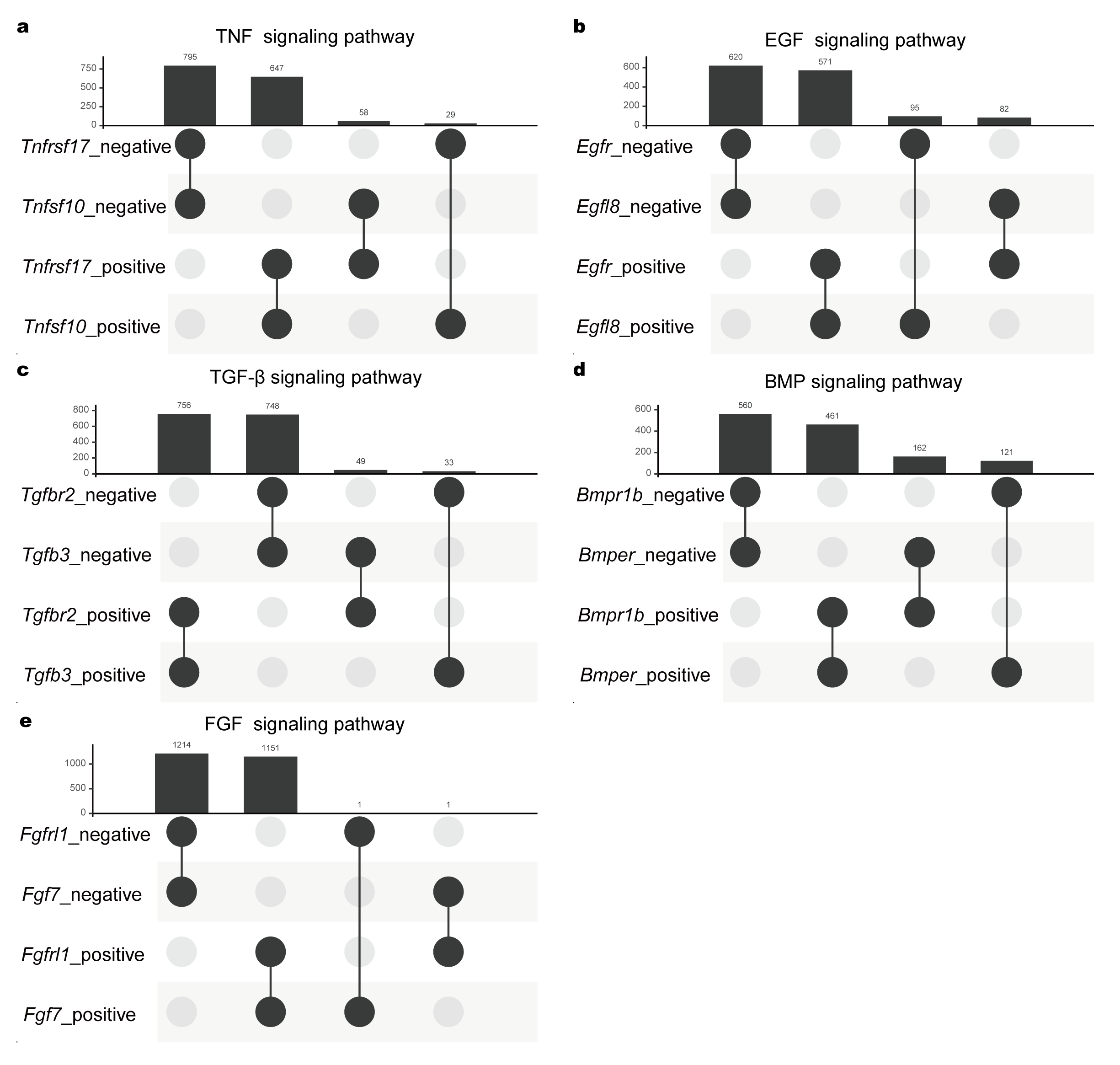

**Supplementary Figure 1. Overlaps of inferred downstream genes for ligand-related and receptor-related genes.**

Upset plots of inferred downstream genes that were positively and negatively regulated by the representative ligand-related and receptor-related genes. (a) TNF signaling pathway; (b) EGF signaling pathway; (c) TGFβ signaling pathway; (d) BMP signaling pathway; (e) FGF signaling pathway.

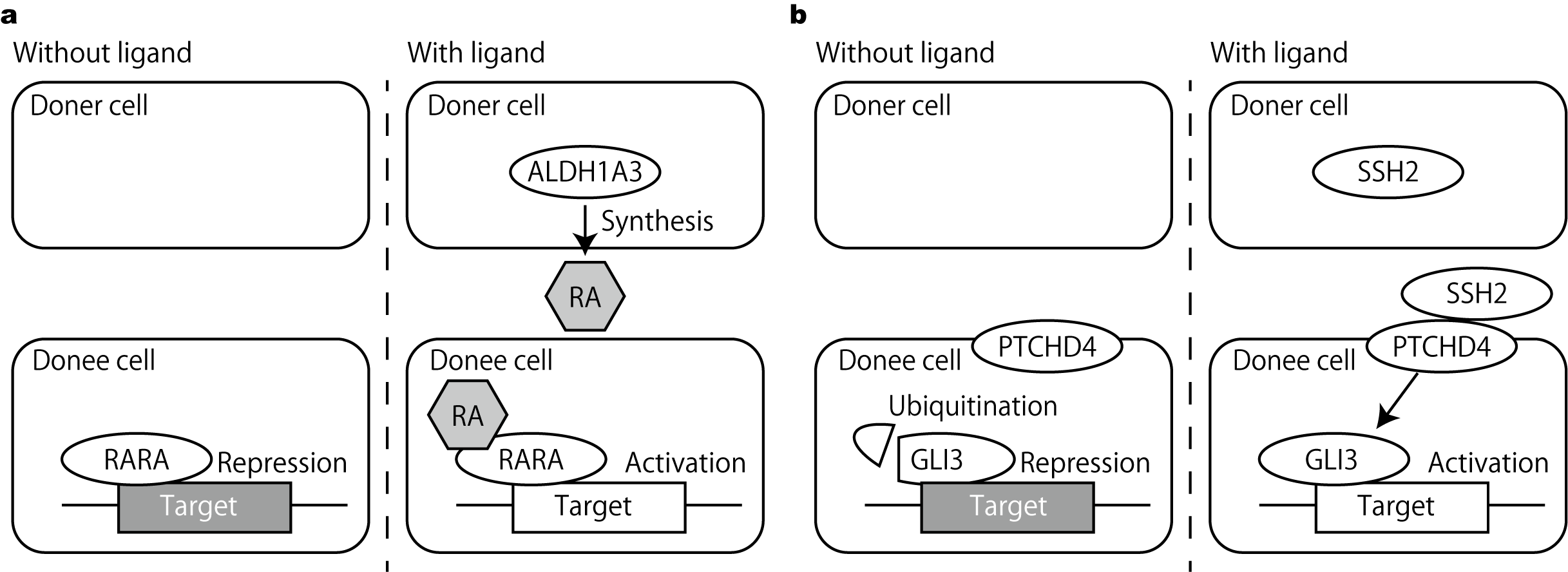

**Supplementary Figure 2. Schematic illustration of the retinoic acid and Hedgehog signaling pathway.**

(a) In the absence of retinoic acid (RA), the RA receptor (RARA) represses target genes. ALDH1A (ALDH1A3) synthesizes RA. In the presence of RA, RA receptor activates the target genes. (b) In the absence of Hedgehog ligand (SSH2), GLI protein (GLI3) is ubiquitinated to function as a repressor of the target genes. In the presence of Hedgehog ligand, PATCHED (PTCHD4) inhibits GLI ubiquitination to activate the target genes.

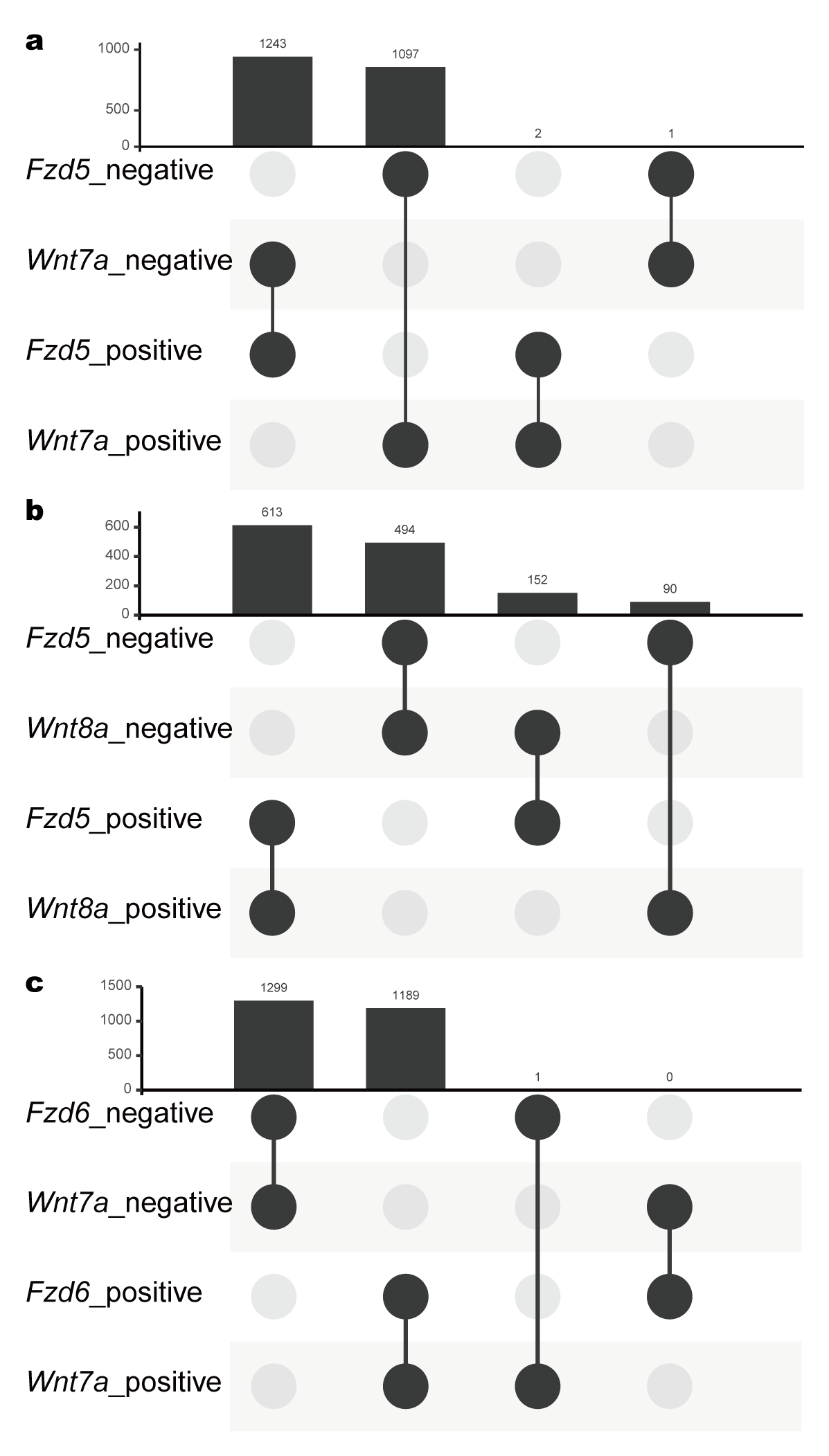

**Supplementary Figure 3. Overlaps of inferred downstream genes of *Wnt7a, Wnt8a, Fzd5,* and *Fzd6.***

Upset plots of inferred downstream genes that were positively and negatively regulated by the representative *Wnt7a* and *Fzd5* (a), *Wnt8a* and *Fzd5* (b), and *Wnt7a* and *Fzd6* (c).

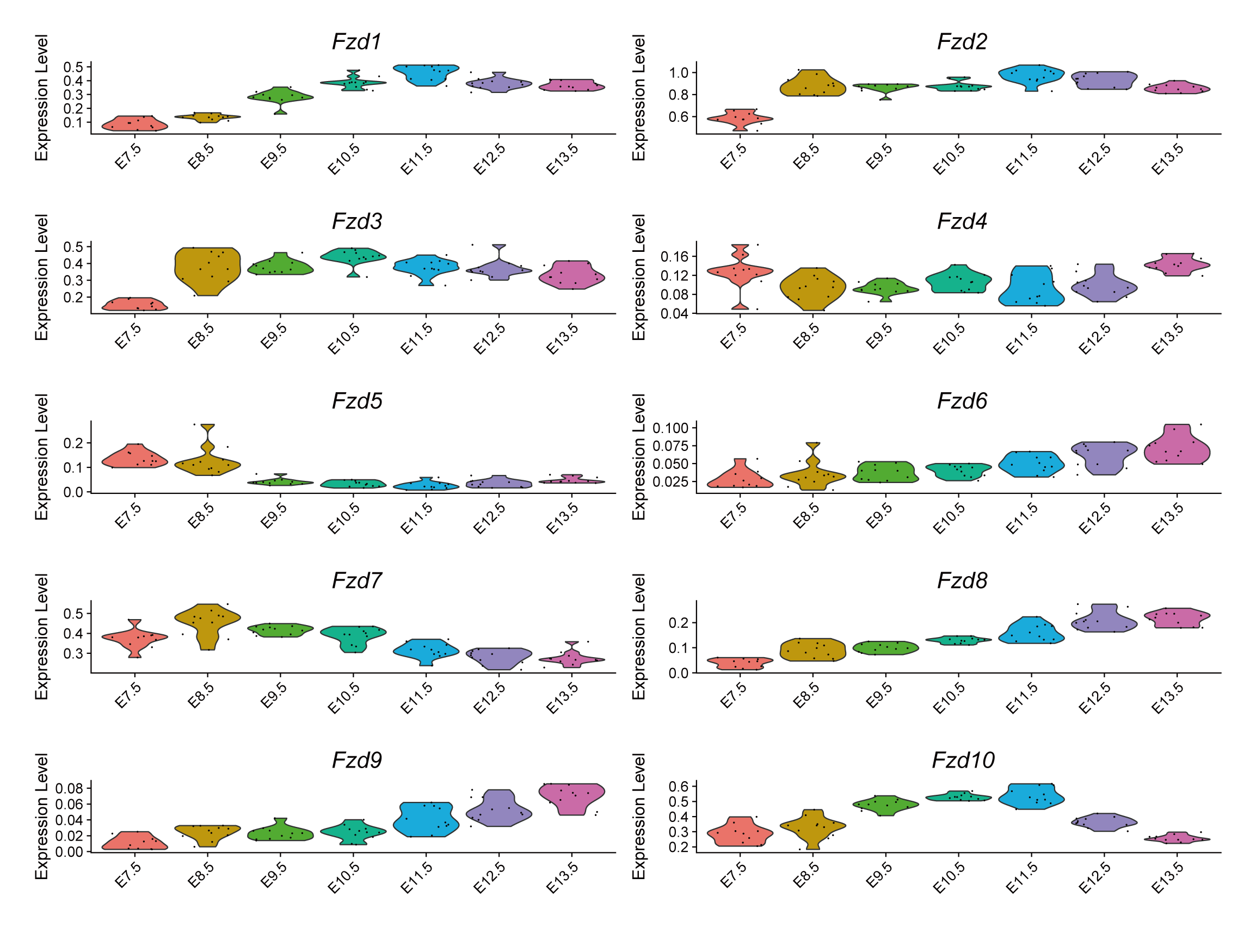
 **Supplementary Figure 4. Gene expression dynamics of *Fzd* genes.**

Violin plots for the relative gene expression of *Fzd* genes from E7.5 to E13.5.

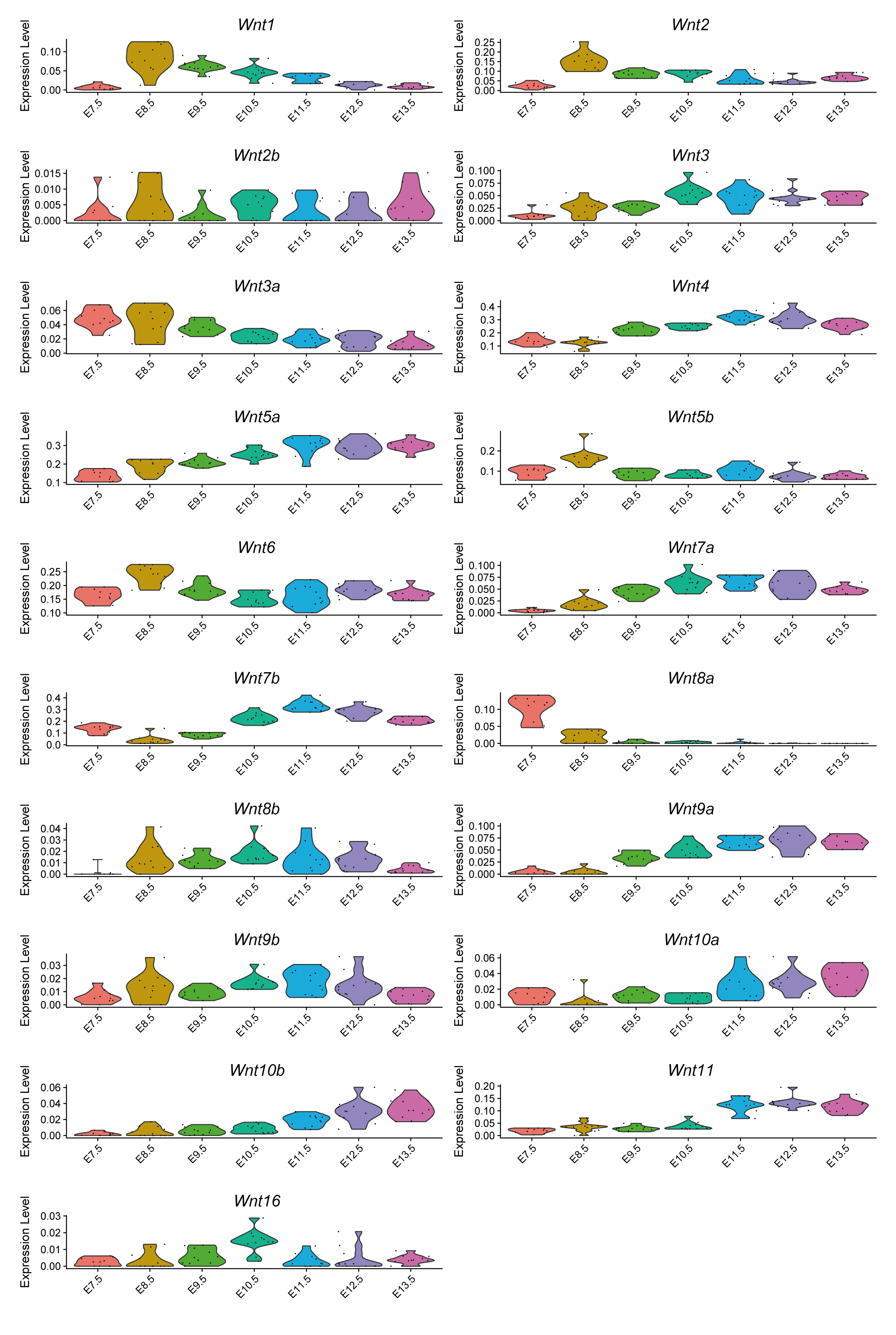

**Supplementary Figure 5. Gene expression dynamics of *Wnt* genes.**

Violin plots for the relative expression of *Wnt* genes from E7.5 to E13.5.
